# Cardiac responses to auditory irregularities reveal hierarchical processing during sleep

**DOI:** 10.1101/2024.09.24.614090

**Authors:** Matthieu Koroma, Paradeisios Alexandros Boulakis, Federico Raimondo, Vaia Gialama, Christine Blume, Mélanie Strauss, Athena Demertzi

**Affiliations:** CRC Human Imaging Unit, GIGA Institue, Allée du 6 Août, 8 (B30), 4000 Sart Tilman, University of Liège, Liège, Belgium; Department of Psychology, Psychology and Neuroscience of Cognition Research Unit, University of Lieège, Liège, Belgium; Institute of Neuroscience and Medicine, Brain and Behaviour (INM-7), Research Center Jülich, Jülich, Germany; Institute of Systems Neuroscience, Medical Faculty, Heinrich Heine University Düsseldorf, Düsseldorf, Germany; Centre for Chronobiology, Psychiatric Hospital of the University of Basel, Basel, Switzerland; Research Cluster Molecular and Cognitive Neurosciences, University of Basel, Basel, Switzerland; Department of Biomedicine, University of Basel, Switzerland; Université libre de Bruxelles (ULB), Hôpital Universitaire de Bruxelles (H.U.B), CUB Hôpital Érasme, Neurology Department and Sleep Unit, Route de Lennik 808, 1070 Brussels, Belgium; Laboratory of Experimental Neurology, ULB Neuroscience Institute, Université Libre de Bruxelles, Brussels, Belgium

**Keywords:** brain-heart interactions, ECG, local-global paradigm, sleep, auditory processing

## Abstract

Our ability to process environmental stimuli varies during sleep. Although much research focused on neural processing, emerging evidence shows that bodily signals may play a key role in understanding the variations of high-level sensory processing. Here, we investigated the cardiac responses to the local-global paradigm, a typical oddball task probing the processing of simple (local) and complex (global) sensory irregularities across unresponsive conscious states. To do so, we analyzed electrocardiography signals in response to auditory deviants obtained during wakefulness and sleep in a total of 56 participants. Our findings revealed that cardiac activity slowed down after global deviants during Rapid-Eye Movement (REM) sleep, especially in tonic REM sleep, *i.e*., in absence of rapid eye movements and further uncovered that cardiac acceleration effects could also be present in non-REM sleep, restricted to light NREM sleep stage, *i.e*, in absence of slow-wave activity. Overall, our findings show that cardiac responses to auditory irregularities can inform the variations of hierarchical information processing across sleep states. They also highlight the embodiment of sleep functions and the value of cardiac signals to study the bodily correlates of sensory processing during sleep.

## Introduction

Sleep is classically considered as a state of loss of vigilance to the external world (Peigneux et al., 2001). Yet, remaining sensitive to the environment during sleep might be crucial for survival, allowing, for example, to wake up if unexpected events occur (Blume & Schabus, 2019; Cirelli & Tononi, 2008; Coenen, 2024). Previous investigations on the neural responses to sounds of different complexities have revealed that the sleeping brain indeed processes from simple acoustic stimuli up to semantic information, but that this ability depends greatly on sleep states (Andrillon & Kouider, 2020; Hennevin et al., 2007).

In comparison, how the body is involved in complex auditory information processing during sleep has been so far understudied (Koroma et al., 2024). Yet, emerging evidence suggests that bodily signals also play a fundamental role in cognition and perception (Azzalini et al., 2019). Empirical support highlighting the importance of bodily signals has been obtained by showing for example that cardiac activity responses to sound inform about the classification of disorders of consciousness beyond neural activity in the local-global paradigm (Raimondo et al., 2017).

The local-global paradigm is a classical mismatch detection task distinguishing between an automatic detection of low-level auditory irregularities (local mismatch effect) and a consciousness-dependent detection of high-level auditory irregularities (global deviance effect) (Bayne et al., 2024; Bekinschtein et al., 2009). By applying this paradigm to healthy sleep, neural signatures of local mismatch detection, but not of global deviance detection, have been previously reported, showing that a loss of vigilance to hierarchical information during sleep (Blume et al., 2022; Strauss et al., 2015, 2022). However, the bodily correlates of hierarchical information processing remained to be investigated in this dataset to probe whether cardiac activity also informs about variations of complex information processing during sleep.

In the present study, we thus tested how cardiac responses to auditory irregularities of different complexities were shaped during wakefulness and sleep. In line with brain results, we hypothesized that local mismatch and global deviance effects would be observed at the cardiac level during wakefulness (Blume et al., 2022; Strauss et al., 2015, 2022). In contrast, only local mismatch effects would be found during sleep, reflecting the breakdown of hierarchical information processing (Blume et al., 2022; Makov et al., 2017; Strauss et al., 2015).

As a secondary hypothesis, we predicted, as observed at the cerebral level, that cardiac effects would be modulated depending on the presence of microphysiological hallmarks of sleep states involved in the gating of sensory processing (Andrillon et al., 2016). Specifically, we hypothesized that, if obtained, cardiac modulations would be observed in light NREM (*i.e*., NREM2) and periods of REM sleep without eye movements (*i.e*., tonic REM), during which a wide range of sensory processing (*e.g*., preferentially reacting to one’s own name) (Andrillon & Kouider, 2020; Perrin, 1999). In contrast, cardiac responses would be absent in deep NREM (*i.e*., NREM3) and periods of REM sleep without eye movements (*i.e*., phasic REM), during which auditory information processing is typically suppressed (Ermis et al., 2010; Koroma et al., 2020; Legendre et al., 2019).

## Methods

### Local-global paradigm during sleep

We extracted the electrocardiographical (ECG) signal of 56 healthy young adults from 18 to 35 years old who participated in the local-global paradigm during wakefulness and sleep (n=28 for a morning nap from Strauss et al., 2015; n=28 for 2 full nights from Blume et al., 2022). Auditory stimuli were pairs of phonetically distant vowels (100 ms duration). Five vowels with a 50 ms interstimulus interval were included in each trial. The first four vowels were identical, and the last one was either the same (“local standard”) or different (“local deviant”; cf. Figure 1A, top). Full details of the protocol are presented in Strauss and colleagues (2015) and Blume and colleagues (2022).

**Figure 1.**
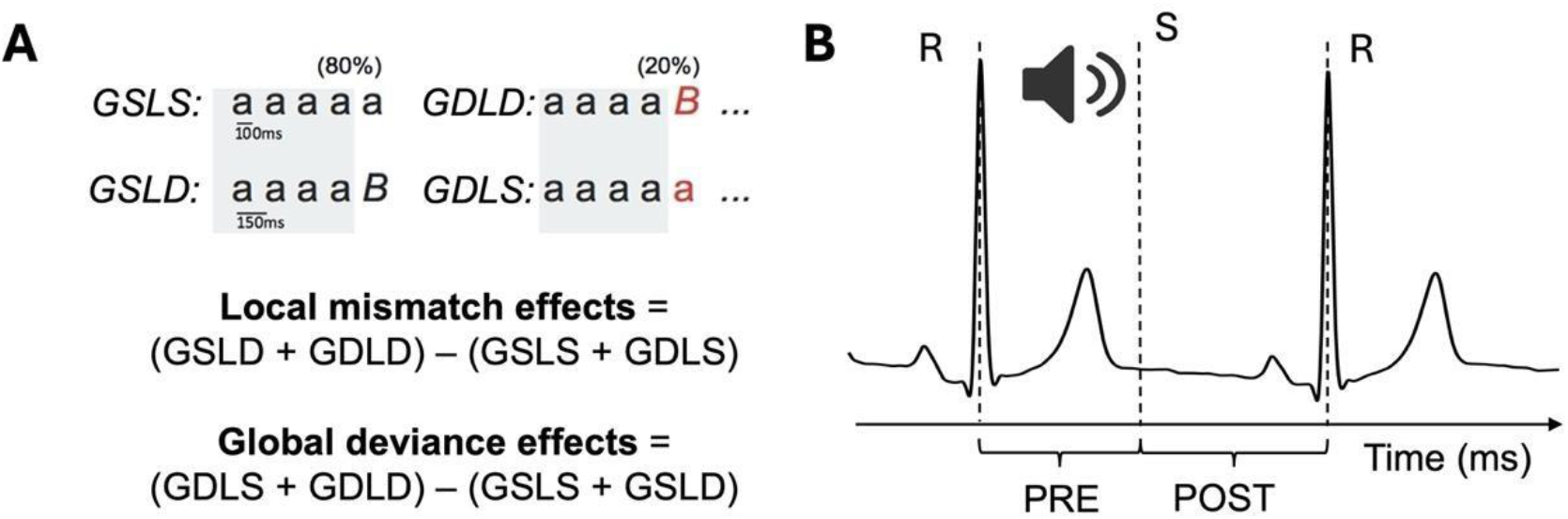
Analysis of the local-global paradigm played during sleep across two datasets. **(A)** A trial was composed of a sequence of 5 tones and belonged to one of the 4 following categories: global standard and local standard (GSLS, top left), global deviant and local deviant (GDLD, top right), global standard and local deviant (GSLD, bottom left), global deviant and local standard (GDLS, bottom right). The combination of responses to these trials allows to define local mismatch and global deviance effects. Adapted from (Strauss et al., 2022). **(B)** Heartbeat was extracted from ECG. For each trial, the duration between onset of the 5th sound with the preceding heartbeat (PRE) and with the following heartbeat (POST) were computed following the methodology of Raimondo and colleagues (2017). Note: S = first sound of the tone sequence; R = cardiac R-peak

Overall, 6914 trials±[1802, 18249] (median ± 95% confidence interval) per participants were presented in sequences of 100 trials in two types of blocks. In the first type of blocks, 80% of the sequences were composed of five identical sounds (aaaaa), referred as “Global Standard Local Standard” (GSLS), and 20% of four identical sounds and a deviant (aaaaB), referred as “Global Deviant Local Deviant” (GDLD). In the second type of blocks, 80% of the sequences were composed of four identical sounds and a deviant (aaaaB), referred as “Global Standard Local Deviant” (GSLD), and 20% of five identical sounds (aaaaa), referred as “Global Deviant Local Standard” (GDLS; Figure 1A, top). The vowel (*e.g*., a or b) serving as standard and deviant was counterbalanced across blocks.

Using the same methodology as adopted for cerebral signals (Bekinschtein et al., 2009; Blume et al., 2022; Strauss et al., 2015), the local mismatch effect was defined by subtracting local deviants from the local standards: (GSLD+GDLD) - (GSLS+GDLS); and the global deviance effect by subtracting global deviants from global standards: (GDLS+GDLD) - (GSLS+GDLD) (Figure 1A, bottom). To further investigate the origin of local mismatch and global deviance effects, analyses were also conducted during sleep at the level of each type of trials presented, *i.e*. GSLS, GSLD, GDLS, and GDLD (Figure S1).

### Cardiac analyses during wakefulness and sleep

Heartbeats were extracted after cleaning the ECG signal (sampled at 120 Hz) using the *ecg_process* function (high-pass Butterworth filter at 0.5 Hz and powerline filter at 50 Hz) and identifying R-peaks with the *neurokit* method from the Neurokit2 package (Makowski et al., 2021). For assessing variations of baseline cardiac activity, interbeat intervals (RR) were computed as a proxy for heart rate during wakefulness and sleep. The values were averaged for wakefulness and each sleep stage within each participant.

Following previous methodology (Raimondo et al., 2017), the cardiac responses to local mismatch and global deviance were obtained by comparing the modulation of the heartbeat timing before and after the occurrence of low-level and high-level auditory irregularities during wakefulness and sleep. The interval between the onset of the fifth sound of each trial with the preceding and following heartbeats defines a PRE and a POST period for each trial (Figure 1C) respectively. To ensure sufficient signal-to-noise ratio, only subjects with more than 10 deviant trials in wakefulness, NREM and REM were retained for analyses.

As in Raimondo and colleagues (2017), our analyses were first restricted to trials for which both PRE and POST intervals were contained between 20 and 600 ms around the fifth sound onset (Figure S1). To account for the fact that NREM is characterized by a lower heart rate than wakefulness (Figure S2A), we also compared PRE and POST periods in wakefulness and sleep using a grid-search approach on time windows ranging from 20 ms to 600 ms, and up to 1200 ms with 100 ms increments, to further explore the presence of local mismatch and global deviation effects in trials with longer interbeat intervals (Table S1).

### Comparisons of cardiac modulation during wakefulness and sleep

Sleep scoring was performed offline according to the AASM guidelines (Blume et al., 2022; Iber et al., 2007; Strauss et al., 2015). Eye movements in REM were identified by independent scorers (M.S. for the Strauss dataset; M.K. and V.G. for the Blume dataset), defining 20±12% (mean±standard deviation) trials in REM as phasic REM, in line with the proportion reported in the literature (Arnulf, 2011; Spreng et al., 1968) (Figure S2B). NREM1 was excluded from analyses as it is considered a transitionary sleep stage (Lacaux et al., 2024; Ogilvie, 2001).

The modulation of heart rate and local mismatch and global deviance effects for PRE and POST periods across sleep stages were tested with linear mixed models with participants as random factor using the *lmer* package in R. Post-hoc comparisons were tested with non-parametric Wilcoxon and Mann-Whitney tests using the *scipy* toolbox and absence of effects with Bayes Factors analyses using the *Pinguoin* toolbox in Python. Multiple comparisons were corrected using false detection rate (alpha threshold=0.05) (Benjamini & Yekutieli, 2001).

Median and 95% confidence interval (95CI) of beta estimates (β), standardized β and t values (t) are reported for mixed-model analyses. Median and 95CI (bootstrap, n=2000) were reported for POST *vs*. PRE comparisons for the grid-search analysis and individual trial analyses. Effect sizes, according to the formula r = z/√n, z being the z-stats of the Wilcoxon signed rank test and n being the number of participants, as well as Bayes Factors (BF) with values above 3 being considered as positive evidence in favor of the null hypothesis (Kass & Raftery, 1995), were also reported for local mismatch and global deviance analyses.

## Results

### Confirmatory: Cardiac activity slows down after global deviants in tonic REM

In terms of the local mismatch effect, there was no evidence for a cardiac modulation (PRE *vs*. POST: β=4.11±[-3.0, 11.23], t(256)=1.14, p=0.26, standardized β=0.24±[-0.17, 0.64]), and the modulation did not differ across vigilance stages (POST *vs*. PRE with: wakefulness *vs*. NREM: β=0.43±[-9.63, 10.49], t(256)=0.08, p=0.93, standardized β=0.02±[-0.55, 0.60]; wakefulness *vs*. REM: β=3.47±[-7.18, 14.11], t(256)=0.64, p=0.52, standardized β=0.20±[- 0.41, 0.81]). Post-hoc testing confirmed the absence of the local mismatch effect in wakefulness, NREM and REM (all tests: p>0.05, BF>3.0) (Figure 2A).

**Figure 2.**
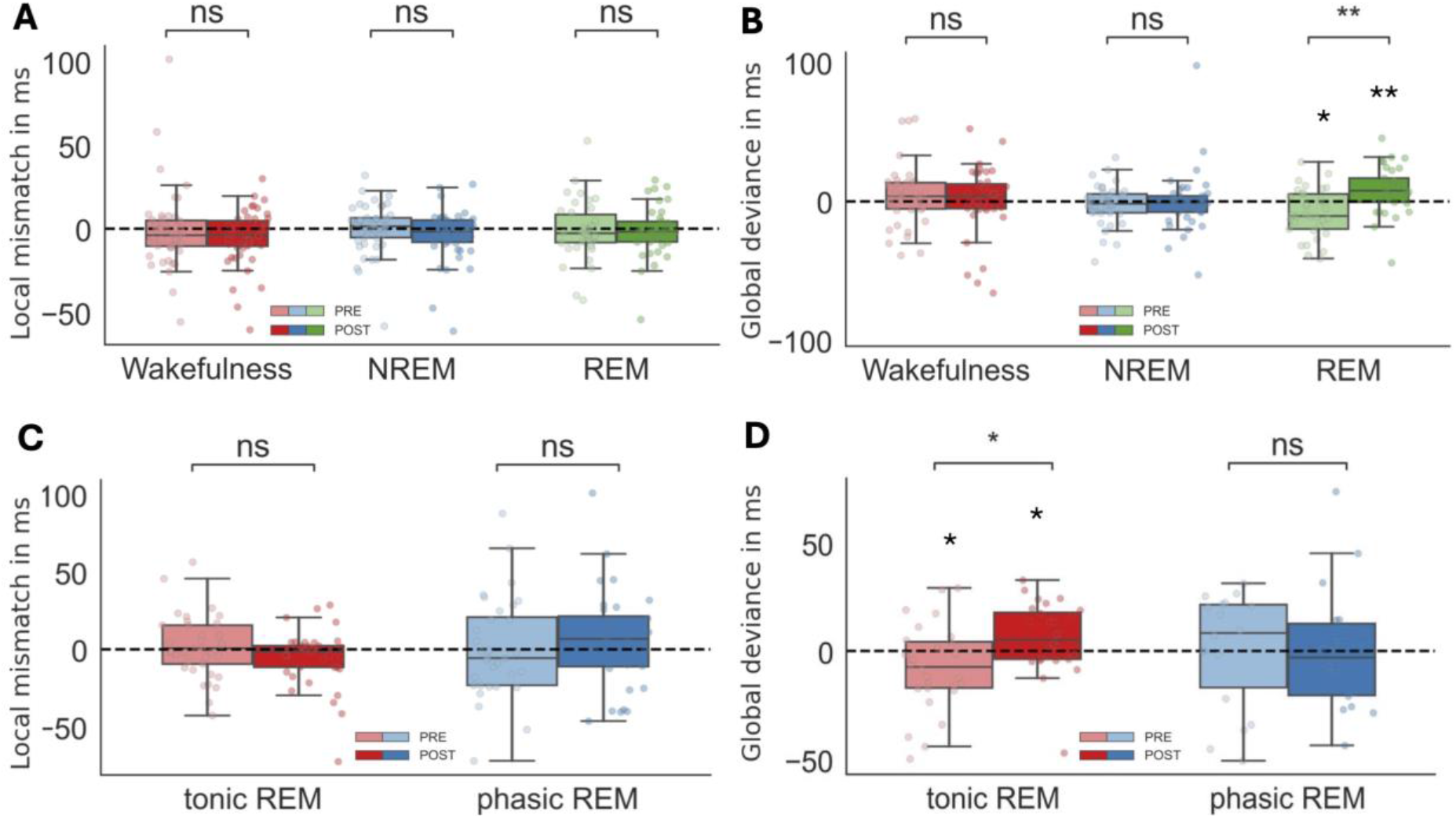
Cardiac activity slows down after global deviants in tonic REM. Local mismatch (**A, C**) and global deviance (**B, D**) effects were computed separately for PRE and POST periods between 20 and 600ms in wakefulness, NREM and REM (top) and tonic REM and phasic REM (bottom). Boxplots represent median, first and fourth quartiles and whiskers deciles for each condition. Datapoints represent individual subjects. Statistical significance for post-hoc test, ns: not significant, *: p<0.05

In terms of the global deviance effect, there was an interaction effect with wakefulness and REM (POST *vs*. PRE and wakefulness *vs*. REM: β=19.91±[6.52, 33.29], t(224)=2.93; p=0.004, standardized β=0.95±[0.31, 1.58]). Post-hoc effects uncovered a cardiac deceleration after global deviants in REM (POST *vs*. PRE: r=-0.52, p=0.002; PRE: -9.37±[-11.95, 1.38], r=-0.39, p=0.02; POST: 8.76±[4.51, 13.72], r=0.54, p=0.004), but not wakefulness nor NREM (all tests: p>0.05, BF>3.0) (Figure 2B).

Post-hoc tests confirmed the absence of a cardiac modulation after local deviants in tonic REM and phasic REM (p>0.05, BF>3.0 for all tests) (Figure 2C). They however showed the presence of a cardiac deceleration after global deviants in tonic REM (POST *vs*. PRE: r=0.19, p=0.02; PRE: r=-0.12, p=0.12; POST: r=0.23, p=0.03) and its absence in phasic REM (POST *vs*. PRE: r=0.08, p=0.77, BF=4.1; PRE: 9.29±[-10.18, 22.05], r=0.01, p=0.83, BF=4.0; POST: -2.01±[-18.37, 12.12], r=-0.10, p=0.83, BF=4).

### Exploratory: Cardiac activity accelerates after local deviants in light NREM

Our grid search analysis revealed of a local mismatch effect between 20 and 900ms in sleep (POST *vs*. PRE: -5.39±[-10.2, 3.0]; p=0.045, corrected Wilcoxon test) while no local mismatch effects were found in wakefulness nor sleep (all tests: p>0.05, Table S1). Post-hoc comparisons confirmed a cardiac acceleration after local deviants in NREM (POST *vs*. PRE: r=-0.33, p=0.017; PRE: 3.92±[1.36, 5.16], r=-0.35, p=0.02; POST: -2.27±[-4.39, -0.35], r=-0.27, p=0.046) and its absence in wakefulness and REM (all tests: p>0.05, BF>3.0) (Figure 3A).

**Figure 3.**
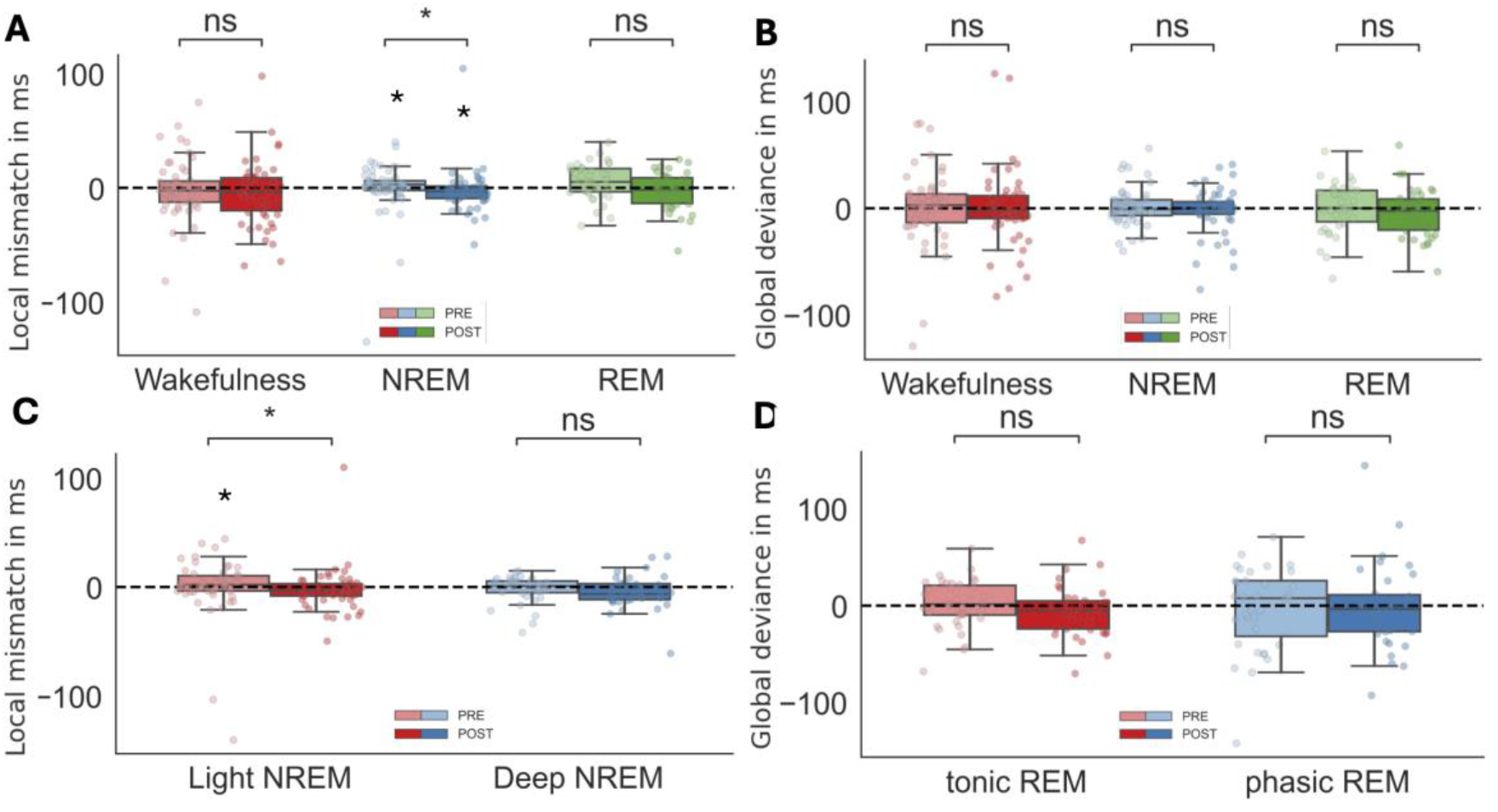
Cardiac activity accelerates after local deviants in light NREM. Local mismatch (**A, C**) and global deviance (**B, D**) effects were computed separately for PRE and POST periods between 20 and 900ms in wakefulness, NREM and REM (top) and light NREM, deep NREM, tonic REM and phasic REM (bottom). Boxplots represent median, first and fourth quartiles and whiskers deciles for each condition. Datapoints represent individual subjects. Statistical significance for post-hoc test, ns: not significant, *: p<0.05

In comparison, our grid-search analysis did not report any global deviance effects between 20 and 900ms (PRE vs POST: -2.7±[-7.7, 2.3]; p=0.497, corrected Wilcoxon test) nor any other global deviance effects in wakefulness nor sleep (all tests: p>0.05) (Table S1). Post-hoc comparisons further confirmed that cardiac activity was not modulated after global deviants in wakefulness, NREM nor REM (all tests: p>0.05, BF>3.0) (Figure 3B).

Finally, post-hoc tests confirmed the presence of local mismatch effects in light NREM (NREM2) (POST *vs*. PRE: r=-0.27, p=0.04; PRE: 2.94±[-1.22, 5.60], r=0.31, p=0.05; POST:-2.56±[-4.23, 0.47], r=-0.22, p=0.10), but not in deep NREM (NREM3) (POST *vs*. PRE: r=-0.24, p=0.16, BF=4.3; PRE: 1.73±[-2.08, 5.34], r=0.08, p=0.65, BF=4.7; POST: -5.43±[-9.15, 1.37], r=-0.28, p=0.18, BF=2.0) (Figure 3C). Post-hoc tests confirmed the absence of global deviance effects in light NREM and deep NREM (all tests: p>0.05, BF>3.0) (Figure 3D).

### Explanatory: Cardiac activity accelerates after local deviants in tonic REM

Finally, we investigated the modulation of cardiac activity in each type of trial separately to precise the origin of local and global effects observed during sleep.

In REM, a cardiac slowdown after global deviants was evidenced in local standard trials (GDLS, POST *vs*. PRE: 17.7±[12.6, 23.1], p<0.001) and local deviant trials (GDLD, POST *vs*. PRE: 13.0±[2.6, 23.1], p=0.034) (Figure 4A). In NREM, local mismatch effects were traced back to the acceleration of cardiac activity after local deviants in global standard (GDLS, POST *vs*. PRE: -2.25±[-4.55, -1.13], p=0.030) but not local standard trials (GDLS, POST *vs*. PRE: - 3.7±[-7.81, 7.16], p=0.326) (Figure 4C).

**Figure 4.**
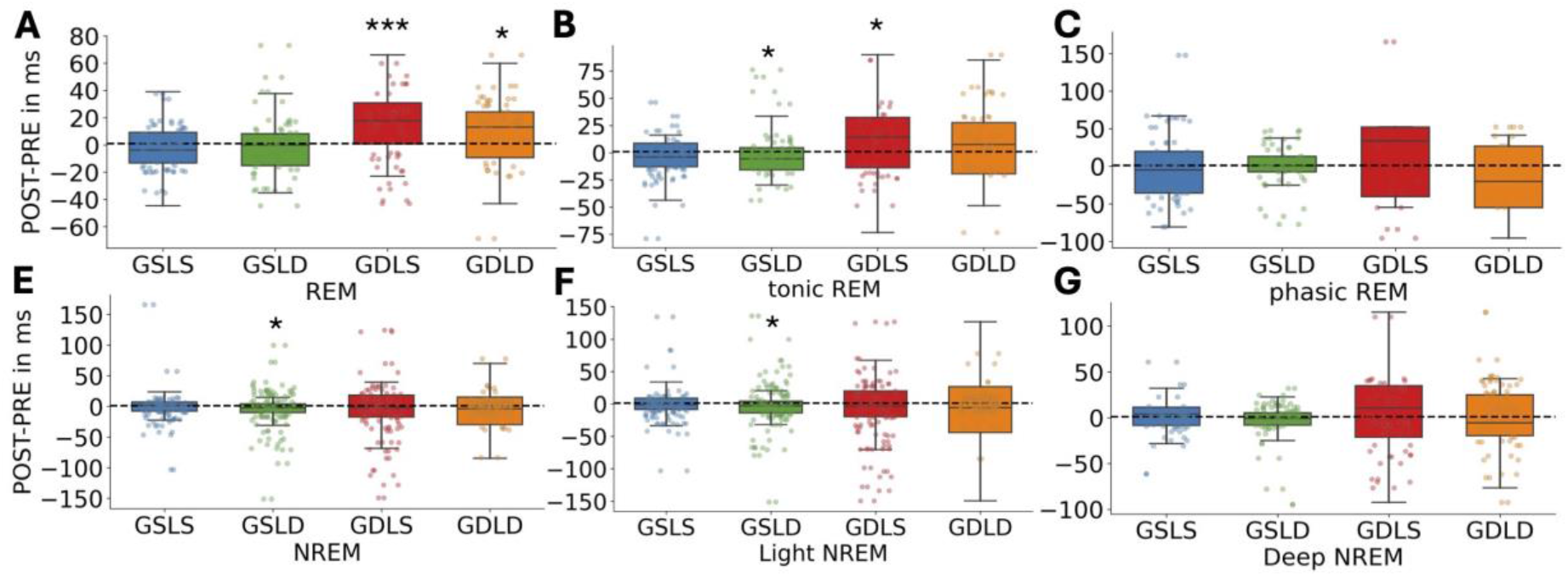
Cardiac activity also accelerates after local deviants in tonic REM. The difference between POST and PRE periods were computed for each type of trial independently after restricting trials between 20 and 600ms for REM (**A**), tonic REM (**B**) and phasic REM (**C**), and between 20 and 900ms for NREM (**D**), light NREM (**E**) and deep NREM (**F**). Boxplots represent median, first and fourth quartiles and whiskers deciles for each condition. Datapoints represent individual subjects. Statistical significance for post-hoc test, ns: not significant, *: p<0.05, ***: p<0.001

In tonic REM, a cardiac slowdown after global deviance was also found in local standard trials (GDLS, POST *vs*. PRE: 14.4±[5.4, 17.1], p=0.025), and a cardiac acceleration was uncovered after local deviants in global standard trials (GSLD, POST *vs*. PRE for GSLD: -5.5± [-13.0, 1.2], p=0.025) (Figure 4B). Comparatively, in phasic REM, no effects were obtained for global deviant in local standard trials (GDLS, POST *vs*. PRE: 33.6±[-40.5, 52.0], p=0.744) and local deviant in global standard trials (GSLD, POST *vs*. PRE: 0.7±[-1.3, 7.3], p=0.569) (Figure 4C).

Finally, in light NREM (NREM2), a cardiac acceleration after local mismatch was confirmed in global standard trials (POST *vs*. PRE, GSLD: -3.1±[-7.8, 0.0], p=0.029), but not in global deviant trials (GDLD: -5.7±[-19.5, 7.5], p=0.126) (Figure 4E). In deep NREM (NREM3), no effects were comparatively found in global standard trials (POST *vs*. PRE, GSLD: -0.9±[-3.8, 1.7], p=0.417) nor in global deviant trials (GDLD, POST vs. PRE: -5.6±[-8.3, 8.6], p=0.732) (Figure 4F).

## Discussion

We investigated the modulation of cardiac activity by hierarchical auditory irregularities presented during wakefulness and sleep. Our results first replicate observations that cardiac responses, contrary to brain responses, are not modulated by auditory irregularities in awake controls (Raimondo et al., 2017). This absence of effect can result from the regulation of cardiac activity by the prefrontal cortex during wakefulness that is deactivated during sleep (Muzur et al., 2002; Wong et al., 2007).

Contrary to our predictions, complex auditory irregularities (global deviance effects) were found in REM sleep, adding novel findings to previous results obtained at the cerebral level (Blume et al., 2022; Strauss et al., 2015). In line with our predictions, we also showed that basic auditory irregularities (local mismatch effect) can also be detected in NREM sleep and that modulations of cardiac activity by auditory stimulation are evidenced in absence of respectively rapid eye movements in REM sleep, *i.e*., tonic REM, and slow-wave activity, *i.e*., light NREM, confirming hallmarks of sleep states as typical markers of reduced sensory processing and their cognitive processing of auditory stimuli (Andrillon & Kouider, 2020; Ermis et al., 2010).

Our findings can be interpreted in the light of the variations of brain-heart interactions during sleep (Chouchou & Desseilles, 2014). In that framework, cardiac slowdown observed after global deviance in tonic REM results from the cerebral processing of sensory information and interoceptive signals at a higher-level (*e.g*., insula) (Bamiou et al., 2003; Oppenheimer & Cechetto, 2016) controlling the release of cardiac sympathetic control (Postuma et al., 2010). Such bradycardia effects are indeed typically associated with error-related monitoring (Skora et al., 2022) or fear-related processing (Battaglia et al., 2024).

Cardiac acceleration after local mismatch in light NREM effects results on their end from the automatic arousal reactions in lower-level brain areas (*e.g*., brainstem) (Vigo et al., 2010; Viola et al., 2011) associated with the release of parasympathetic tone (Benarroch, 2018; Griffiths et al., 2001). Such tachycardia effects align well with previous reports of cardiac accelerations associated with transient arousal increases that are observed upon the presentation of auditory mismatch in NREM sleep even in absence of detectable EEG acceleration or cortical arousal effects (de Zambotti et al., 2018).

Our results further extend to non-pathological cases previous findings showing that, in disorders of consciousness patients, cardiac responses to the auditory local-global paradigm provide additional information to the classification of conscious states beyond cerebral signals (Raimondo et al., 2017). In that study, global deviance effects were observed in minimally conscious patients who exhibit more complex behaviors and cognitive than unresponsive wakefulness syndrome patients (Laureys, 2005). This confirms our findings that the effects of global deviation observed in REM sleep are also present in states characterized by increased complexity of cognitive processing during sleep (Tononi et al., 2024; Tononi & Massimini, 2008).

Our methodology complements the study of Raimondo and colleagues (2017) by showing that exploring PRE and POST periods over different time windows as well as reporting responses to the different types of trials provide additional insights on the modulation of cardiac activity by auditory stimulation. We thus demonstrate that analyzing the dynamics of cardiac responses to sounds during sleep on extended time windows and trial types is also relevant for uncovering correlates of sensory processing across conscious states.

Future studies might also investigate the cerebral processing of cardiac signals (*e.g*., heart-evoked potential) to understand how cardiac activity also influences in return brain responses and thus offer an integrated picture of the brain-body correlates of auditory irregularity processing during sleep, as it has been obtained for disorders of consciousness patients (Candia-Rivera et al., 2023).

In conclusion, we demonstrate with this study that investigating the cardiac correlates of the local-global paradigm during sleep reveal the preservation of the processing of complex irregularities, *i.e*., global deviance effects, in REM sleep and of basic irregularities, *i.e*., local mismatch effects, in NREM sleep. These effects are observed in absence of microphysiological markers of sleep disconnection, respectively rapid eye movements in REM sleep and slow-wave activity in NREM sleep. Some of our findings crucially diverge from our initial predictions based on cerebral signals (Blume et al., 2022; Strauss et al., 2015, 2022), adding to previous demonstrations that cardiac signals encode additional information about sensory processing as compared to brain activity alone (Koroma et al., 2024; Whitehurst et al., 2016, 2022). Overall, our study provides novel evidence supporting the embodiment of cognitive functions and highlights the value of cardiac signals to identify the correlates of complex information processing across conscious states.

## Data and code availability

Codes and dataframes used for analyses are in open access via https://gitlab.uliege.be/poc/sleeplocalglobal

## Author contributions

### Matthieu Koroma

Conceptualization; investigation; writing – original draft; methodology; validation; visualization; writing – review and editing; project administration; formal analysis; software; data curation; resources; funding acquisition; supervision.

### Paradeisos Boulakis

Methodology; validation; writing – review and editing; resources.

### Vaia Gialama

Methodology; validation; formal analysis.

### Federico Raimondo

Conceptualization; methodology; validation; writing – review and editing; resources.

### Christine Blume

Methodology; validation; visualization; writing – review and editing; project administration; formal analysis; data curation; resources; supervision.

### Mélanie Strauss

Conceptualization; Methodology; validation; visualization; writing – review and editing; project administration; formal analysis; data curation; resources; supervision; funding acquisition.

### Athena Demertzi

Supervision; conceptualization; investigation; validation; visualization; writing – review and editing; formal analysis; project administration; resources; data curation; methodology; writing – original draft; funding acquisition.

