## Supplementary Information for "Cardiac responses to auditory irregularities reveal hierarchical processing during sleep"

**Supplementary Figures**

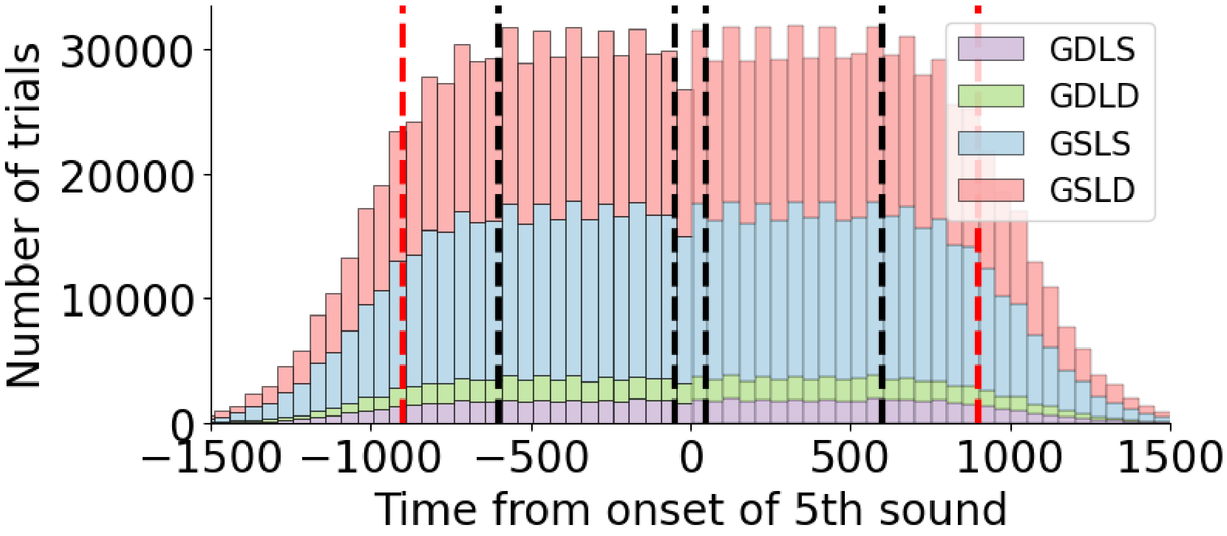


Figure S1. **PRE and POST periods were defined based on heartbeats around 5th sound onset**. PRE periods were defined based on heartbeats before the onset of the 5th sound (negative values) and POST periods based on heartbeats after the onset of the 5th sound (positive values). GDLS: Global Deviant Local Standard, GDLD: Global Deviant Local Standard, GSLS: Global Standard Local Standard, GSLD: Global Standard Local Deviant. PRE and POST periods were restricted between 20 and 600ms (black dotted lines) and 900ms (red dotted lines).

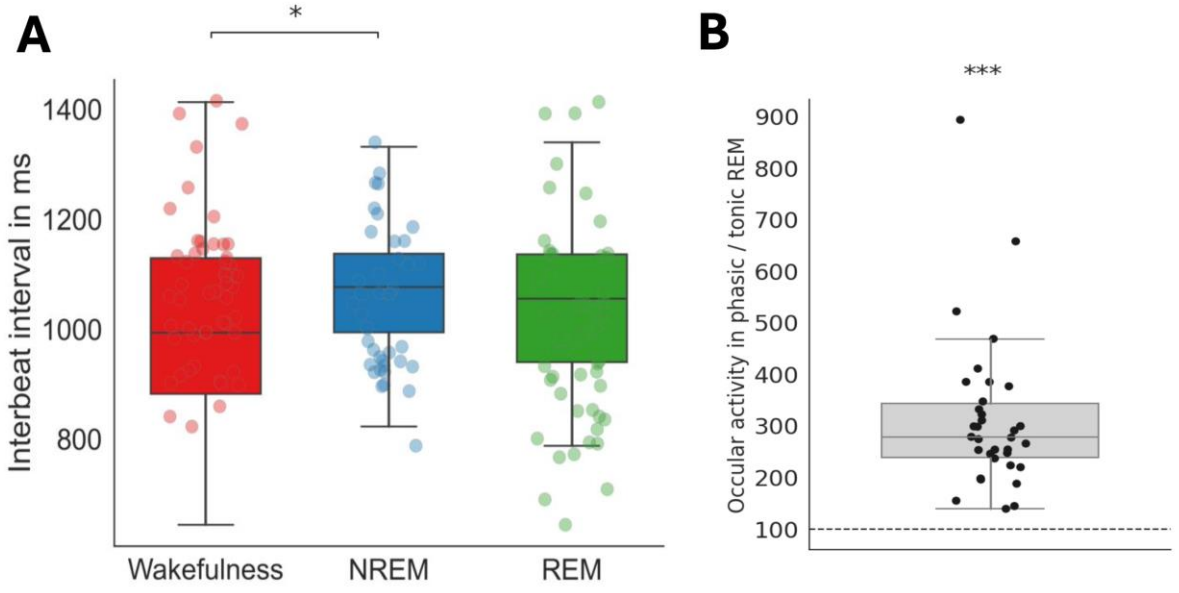


Figure S2. **Cardiac and ocular signals are modulated across sleep stages. (A)** Interbeat interval were computed by extracting R-peaks from electrocardiographic signals and averaging over sleep scoring windows of the same stage for each subject.**(B)** Ocular activity was defined as the standard deviation of electrooculographic signals around trial onset (-4s to 4s). Statistical significance for post-hoc tests, *: $p<0.05$, ***: $p<0.001$

Table S1. **Grid-search analysis approach identifies a local mismatch effect during sleep when including PRE and POST periods from 20 to 900ms**. The median and 95% confidence interval (bootstrap, n=2000) of the POST-PRE and the post-hoc Wilcoxon test POST *vs.* PRE corrected for multiple comparisons were computed for the local mismatch and global deviance effects in wakefulness and sleep (NREM2, NREM3 and REM grouped together) on PRE and POST periods ranging between 20ms and 600 until 1200ms with 100ms step interval. Statistical significance, *: p<0.05

| Effect | State | 600ms | 700ms | 800ms | 900ms | 1000ms | 1100ms | 1200ms |
|---|---|---|---|---|---|---|---|---|
| Local mismatch | Wake | -2.6± [-8.6, 4.9] | -6.3± [-14.1, -1.8] | -3.2± [-8.5, 7.2] | 0.01± [-12.4, 8.4] | -7.3± [-20.3, 0.3] | -3.7± [-16.6, -0.5] | -2.3± [-20.7, -2.5] |
| | Sleep | -2.2± [-5.0, -1.1] | -3.0± [-6.4, -0.8] | -3.4± [-6.0, -0.2] | **-5.4± [-10.2, -3.0]*** | -3.3± [-9.1, 0.9] | -4.3± [-9.8, 0.5] | -4.4± [-9.2, 2.6] |
| Global deviance | Wake | 0.9± [-10.3, 9.1] | -9.3± [-16.1, 4.7] | -2.5± [-12.3, 9.6] | -7.4± [-12.9, 7.9] | -0.1± [-15.4, 17.9] | 1.7± [-13.4, 18.8] | 6.5± [-8.6, 19.4] |
| | Sleep | 3.5± [-1.1, 15.8] | 1.2± [-6.4, 6.1] | 0.8± [-8.8, 7.0] | -2.7± [-7.7, 2.3] | -2.8± [-9.1, 7.4] | -5.6± [-8.3, 3.4] | -4.7± [-9.0, 3.6] |
